# *Pseudomonas syringae* pv. tomato DC3000 Effector HopG1 is a multi-faceted protein that Triggers Necrotic Cell Death that is attenuated by the Nonhost Resistance 2B (AtNHR2B) Protein

**DOI:** 10.1101/2021.11.02.466876

**Authors:** Catalina Rodríguez-Puerto, Rupak Chakraborty, Raksha Singh, Perla Rocha-Loyola, Clemencia M. Rojas

## Abstract

The plant pathogenic bacterium Pseudomonas syringae pv. tomato DC3000 (Pst DC3000) has become a paradigm in plant-bacteria interactions due to its ability to cause disease in the model plant Arabidopsis thaliana. Pst DC3000 uses the type III secretion system to deliver type III secreted effectors (T3SEs) directly into the plant cytoplasm. Pst DC3000 T3SEs contribute to pathogenicity by suppressing plant defense responses and targeting plant’s physiological processes. Although the complete repertoire of effectors encoded in the Pst DC3000 genome have been identified, the specific function for most of them remains to be elucidated. The mitochondrial-localized T3E HopG1, suppresses plant defense responses and promotes the development of disease symptoms. Here, we show that HopG1 triggers necrotic cell death that enables the growth of non-adapted pathogens. We further showed that HopG1 interacts with the plant immunity-related protein AtNHR2B and that AtNHR2B attenuates HopG1-virulence functions.

## Introduction

*Pseudomonas syringae* is a plant pathogenic Gram-negative bacterium that causes diseases in a wide range of plants. Due to this broad host range, the species has been divided into more than 50 pathovars (pv), each pathovar designation based on their host of isolation (1). Among those pathovars, *Pseudomonas syringae* pv. tomato DC3000 (*Pst* DC3000), the causal agent of bacterial speck on tomato, has become a model pathogen to understand bacterial pathogenicity towards plants because it also causes disease in the model plants *Arabidopsis thaliana* and *Nicotiana benthamiana* (1–4). The pathogenicity of *Pst* DC3000 is mostly due to the type III secretion system (T3SS), a complex of proteins encoded by the Hypersensitive Response and Pathogenicity/ Hypersensitive Response and Conserved (*Hrp/Hrc*) genes (5). *Hrp/Hrc*-encoded proteins assemble an apparatus spanning the inner and outer bacterial membranes that enables the bacterium to deliver bacterial proteins (effectors), directly into the host cytoplasm (6–8).

The genome of *Pst* DC3000 encodes 28 type III effectors (T3Es) that are delivered into plant cells (9–11), where acting together interfere with plant immune responses to enable bacterial parasitism (12–14). The plant immune responses include two main branches: 1) Pathogen-Associated Molecular Patterns-Triggered Immunity (PTI) that recognizes conserved features in pathogens known as Pathogen-Associated Molecular Patterns (PAMPs) through surface-localized Pattern Recognition Receptors (PRRs) (15–17), and 2) Effector-Triggered Immunity (ETI), that recognizes pathogen effector molecules by R (resistance) proteins (15, 16, 18). An outcome of ETI is the elicitation of the hypersensitive response (HR), a type of localized programmed cell death, that restricts pathogen proliferation and tissue damage (19–21).

*Pst* DC3000 T3E are functionally redundant, as mutations in individual effectors do not have a significant impact on pathogenicity, yet all the effectors are required for full pathogenicity in tomato and *Arabidopsis* (22). Interestingly, deletion of the T3E HopQ1-1 expands the host range of *Pst* DC3000 to *N. benthamiana* (4). Remarkably, only eight T3Es effectors: AvrPtoB, HopM1, AvrE, HopE1, HopG1, HopAM1-1, HopAA1-1 and HopN1 make up the minimal functional needed for *Pst* DC3000 to be virulent in *N. benthamiana* (23). Among those effectors, HopG1 was previously implicated in the suppression of PTI and ETI responses (24–26). Further research showed that HopG1 localizes to mitochondria (24), where it interacts with the mitochondrial-localized kinesin motor protein to modulate actin cytoskeleton and promote the development of chlorotic symptoms (27).

We previously identified AtNHR2B (*Arabidopsis thaliana* nonhost resistance 2B) as a protein that functions in plant immunity (28–31). *Atnhr2b* mutant plants are immunocompromised, and as a result, they are susceptible to pathogens that normally do not cause disease in wild-type plants, called non-adapted pathogens (31). We further showed by live-cell imaging that AtNHR2B tagged with the green fluorescent protein (GFP) localizes to chloroplasts and compartments of the endomembrane system (31), and through proteomics approaches, we also showed that AtNHR2B interacts with proteins localized to chloroplasts, mitochondria and nucleus (32). Until now, it was not known if AtNHR2B was targeted by pathogen virulence factors.

Here, we show that the *Pst* DC3000 T3E HopG1 interacts with AtNHR2B and that AtNHR2B interferes with HopG1 virulence functions. We also expand the range of functions for HopG1 with the discovery that transient or stable expression of *HopG1* in *N. benthamiana* and Arabidopsis, respectively, triggers necrotic cell death that enables the growth of non-adapted pathogens. Our results and the results of others highlight the multifaceted nature of HopG1 and its interplay with AtNHR2B.

## Results

### Adapted pathogens changed the pattern of localization and abundance of AtNHR2B-GFP

We previously showed that *AtNHR2B* is induced by pathogens and pathogen-derived elicitors, and that *AtNHR2B-GFP* transiently expressed in *N. benthamiana* localizes to the cytoplasm and to highly dynamic structures (punctae) reminiscent of subcellular compartments of the endomembrane system (31). *AtNHR2B* is induced by non-adapted bacterial pathogens (that are unable to cause disease in a particular host), as well as, adapted bacterial pathogens (that cause disease in a specific host), however, we do not know if pathogens’ lifestyle (non-adapted vs adapted) have an effect on the function of AtNHR2B. Since AtNHR2B is an immune-related protein, we hypothesized that adapted pathogens could target this protein for parasitism. To start testing that hypothesis, we evaluated the localization of AtNHR2B-GFP by transient expression in *N. benthamiana. N. benthamiana* plants transiently expressing *AtNHR2B-GFP* were mock-treated with water or inoculated with *Pstab* (adapted pathogen of *N. benthamiana*), or with *Pst* DC3000 (non-adapted pathogen of *N. benthamiana*). Mock-treated plants and plants inoculated with *Pst* DC3000 showed the expected localization to cytoplasm and punctae. Interestingly, after inoculation with *Pstab* (adapted pathogen of *N. benthamiana*), the signal from AtNHR2B-GFP was limited to the cytoplasm only (Fig. 1A).

**Fig. 1.**
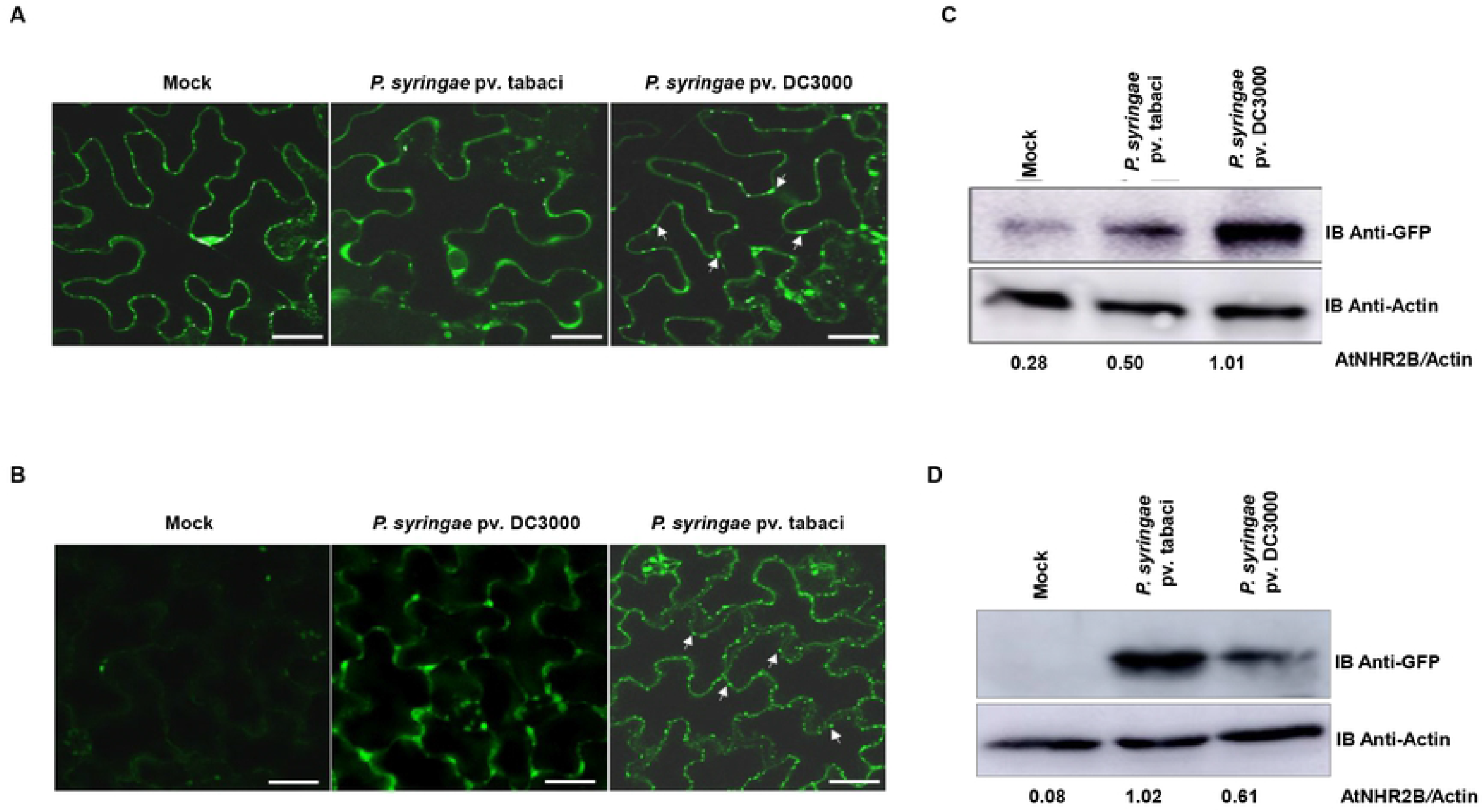
Adapted pathogens interfere with AtNHR2B-GFP localization and protein abundance. (**A**)The host adapted pathogen *Pstab* alters AtNHR2B-GFP localization. *N. benthamiana* plants transiently expressing *AtNHR2B-GFP* were infiltrated with water (mock) or inoculated with either the adapted pathogen *Pstab* or the non-adapted pathogen *Pst* DC3000. Treated leaf samples were collected after 24 hpi and imaged by laser scanning confocal microscopy. Confocal images were taken at 20X magnification and using an excitation and emission wavelengths of 488 nm and 500 to 535 nm, respectively. White arrows show punctate bodies. Scale bar = 10 μm. (**B**) The host adapted pathogen *Pst* DC3000 alters AtNHR2B-GFP localization in *Arabidopsis. Arabidopsis* transgenic plants expressing *AtNHR2B-GFP* were mock-treated or inoculated with the adapted pathogen *Pst* DC3000 and the non-adapted pathogen *Pstab*. Treated leaf samples were collected after 24 hpi and imaged by laser scanning confocal microscopy. Confocal images were taken at 20X magnification and using an excitation and emission wavelengths of 488 nm and 500 to 535 nm, respectively. White arrows show punctate bodies. Scale bar = 10 μm. (**C**) The host adapted pathogen *Pstab* alters AtNHR2B-GFP protein abundance in tobacco plants. *N. benthamiana* plants transiently expressing *AtNHR2B-GFP* were infiltrated with water (mock) or inoculated with either the adapted pathogen *Pstab* and the non-adapted pathogen *Pst* DC3000. Treated leaf samples were collected after 24 hpi to evaluate protein abundance by Western blot using anti-GFP and anti-actin antibodies. Protein quantification was done by ImageJ software (http://rsb.info.nih.gov/ij/), using actin signal for normalization. Number below the protein bands represent the AtNHR2-GFP/Actin ratios. (**D**) The host adapted pathogen *Pst* DC3000 alters AtNHR2B-GFP protein abundance in *Arabidopsis* plants. *A. thaliana* transgenic plants expressing *AtNHR2B-GFP* were infiltrated with water (mock), or inoculated with either the adapted pathogen *Pst* DC3000 and the non-adapted pathogen *Pstab*. Treated leaf samples were collected after 24 hpi to quantify protein abundance by Western blot using anti-GFP and anti-Actin antibodies. Protein quantification was done by ImageJ software (http://rsb.info.nih.gov/ij/), using Actin signal for normalization. Number below the protein bands represent the AtNHR2-GFP/Actin ratios.

The change in AtNHR2B-GFP pattern of localization in *N. benthamiana* associated with inoculation with the adapted pathogen was validated in transgenic *Arabidopsis* plants expressing *AtNHR2B-GFP* from its native promoter (31). In such plants, the fluorescent signal was very low in mock-treated plants, but significantly increased upon inoculation with *Pst* DC3000 and *Pstab*. Similar to the results obtained in *N. benthamiana*, the pattern of AtNHR2B-GFP localization differed depending on the pathogen used for the inoculation; inoculation with the non-adapted pathogen of *A. thaliana, Pstab*, caused the localization of AtNHR2B-GFP to cytoplasm and small punctae as previously described (31) (Fig. 1B). However, inoculation with the adapted pathogen of *A. thaliana*, *Pst* DC3000, changed the pattern of AtNHR2B-GFP localization and the fluorescence signal was detected in the cytoplasm only and appeared diffused and distorted (Fig. 1B).

To evaluate if the diffusion of the fluorescence signal was due to adapted pathogens interfering with protein abundance, we collected leaves from *N. benthamiana* and *Arabidopsis* plants expressing *AtNHR2B-GFP* that were mock-treated, or inoculated with their corresponding adapted and non-adapted pathogens. Collected leaves were used for protein extraction and quantitative Western blot analysis using anti-GFP antibodies to detect AtNHR2B-GFP and anti-actin antibodies for normalization. *N. benthamiana* expressing *AtNHR2B-GFP* and mock-treated, showed low levels of AtNHR2B-GFP. However, inoculation with both adapted (*Pstab*) and non-adapted (*Pst* DC3000) pathogens increased AtNHR2B-GFP concentration in comparison with mock-treated plants, but the amount of AtNHR2B-GFP was reduced by 50% in plant samples treated with *Pstab* in comparison with plants treated with *Pst* DC3000 (Fig. 1C). In a similar way, AtNHR*2B-GFP* transgenic *Arabidopsis* lines did not accumulate any protein after mock treatment and showed ~60% less AtNHR2B-GFP after inoculation with the adapted pathogen *Pst* DC3000 than plants inoculated with the *Pstab* (Fig. 1D). These results demonstrate that adapted pathogens reduce the abundance of AtNHR2B-GFP. Altogether, these results revealed that adapted pathogens alter AtNHR2B-GFP protein localization and abundance.

### The *Pst* DC3000 mutant lacking HopG1 is able to grow better in the *Atnhr2b* mutant background

The finding that the localization and protein abundance of AtNHR2B is altered by adapted pathogens, prompted us to investigate whether AtNHR2B could be a target for *Pst* DC3000 T3E. The finding that HopG1 localizes to mitochondria (24, 27), together with our previous result that AtNHR2B interacts with proteins localized to mitochondria (23), led us to hypothesize that HopG1 could target AtNHR2B. To test that hypothesis, we initially evaluated the growth of wild-type *Pst* DC3000 and the *Pst* DC3000 Δ*hopG1* mutant in wild-type Col-0 and in *Atnhr2b* mutant plants at 3 days post-inoculation (dpi) (Figure 2). As an adapted pathogen of *Arabidopsis*, the wild-type strain *Pst* DC3000 is able to grow in wild-type Col-0 to 10^7^ CFU/cm^2^ at 3 dpi. In contrast, the growth of the *Pst* DC3000Δ*hopG1* was ~10-fold lower than the growth of wild-type *Pst* DC3000. In the *Atnhr2b* mutant plants, the growth of the wild-type *Pst* DC3000 was equivalent to its growth in wild-type Col-0 plants. Interestingly, the *Pst* DC3000Δ*hopG1* mutant grew to higher levels (~10^7^ CFU/cm^2^) in *Atnhr2b* mutant plants and those levels were equivalent to the growth of *Pst* DC3000 in wild-type Col-0. These results suggest that the *Atnhr2b* mutation restores the growth defect in the *PstDC3000*Δ*hopG1* mutant supporting a functional relationship between HopG1 and AtNHR2B.

**Figure 2.**
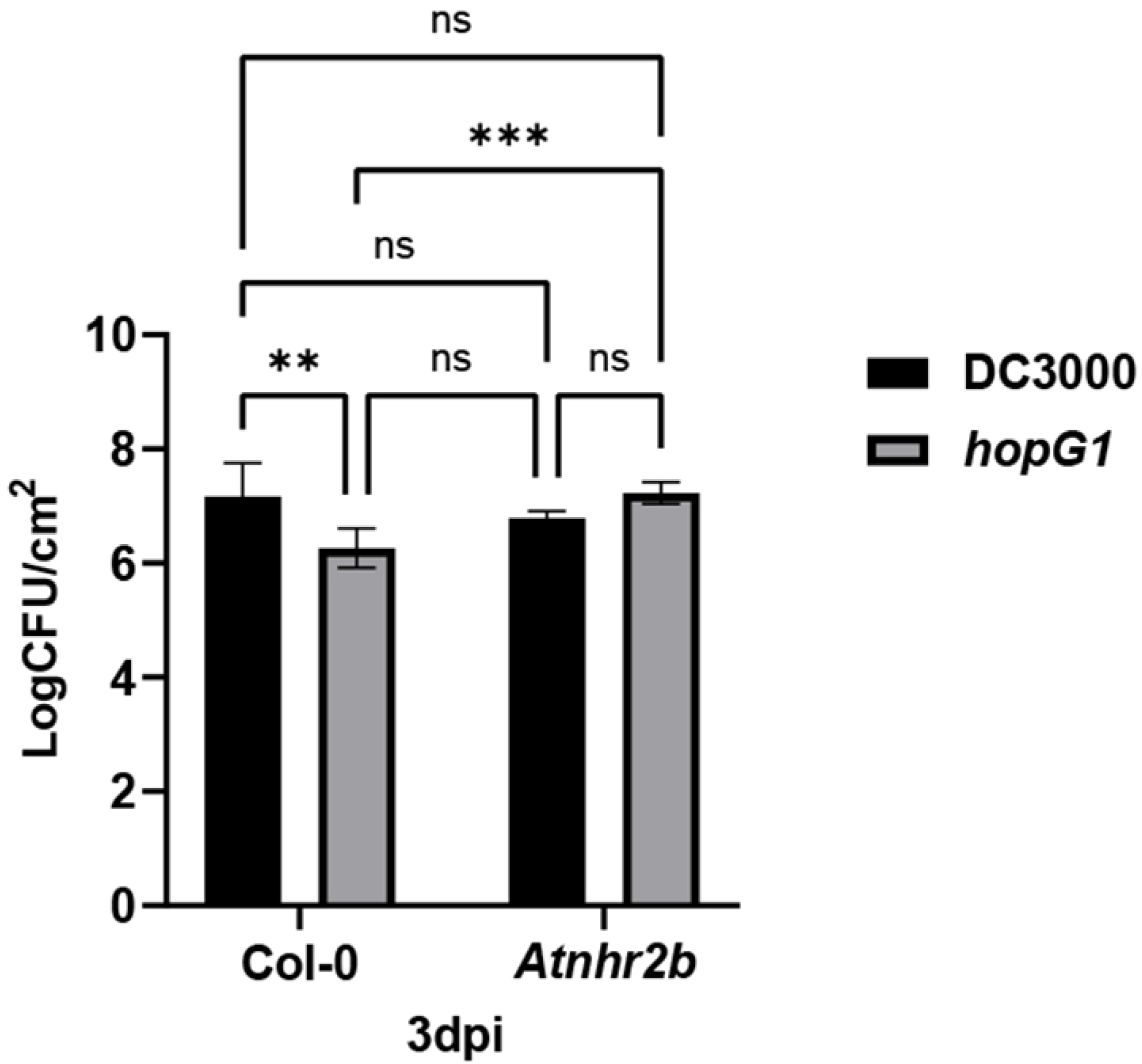
The growth defect in *Pst* DC3000 *hopG1* deletion mutant is restored in *Atnhr2b* mutant *Arabidopsis* plants. Wild-type Col-0 and the *Atnhr2b* mutant were syringe-inoculated with the wild-type *Pst DC3000* and the *Pst* DC3000 Δ *hopG1* mutant at a concentration of 1 × 10^6^ CFU/ml. Inoculated leaves were collected at 3 dpi to enumerate bacterial populations. Bars represent means and standard deviation for bacterial growth in Col-0 (white bars) and *Atnhr2b* (black bars) for four replications. Asterisks above the bars represent statistically significant difference in the growth of wild-type *Pst DC3000* and the *Pst* DC3000 Δ *hopG1* in Col-0 versus their respective growth in the *Atnhr2b* mutant plants *at* 3 dpi using a two-way ANOVA and Sidak’s multiple comparison test with P-value ≤0.05. Number of asterisks reflects levels of significance, with ns equivalent to P≥0.05, * to P≤0.05, ** to P≤0.01, *** to P≤0.001 and **** to P≤0.0001. The experiments were repeated three times with consistent results.

### The *Pst* DC3000 Effector HopG1 Interacts with AtNHR2B

To start dissecting the functional relationship between HopG1 and AtNHR2B, we tested the physical interaction between AtNHR2B and HopG1 with the yeast two-hybrid system using HopG1 as a bait and AtNHR2B as a prey. Yeast co-transformed with *pDEST32::HopG1* and *pDEST22::AtNHR2B* grew on Triple Drop Out (TDO) media (-leu,-trp,-his) containing 15mM 3-AT indicating the transcriptional activation of histidine biosynthetic genes as a result of the interaction between HopG1 and AtNHR2B. That interaction is not the result of autoactivation, because yeast transformed with the empty vector *pDEST32* and *pDEST22::AtNHR2B* did not grow on TDO + 15mM 3-AT (Figure 3A).

**Figure 3.**
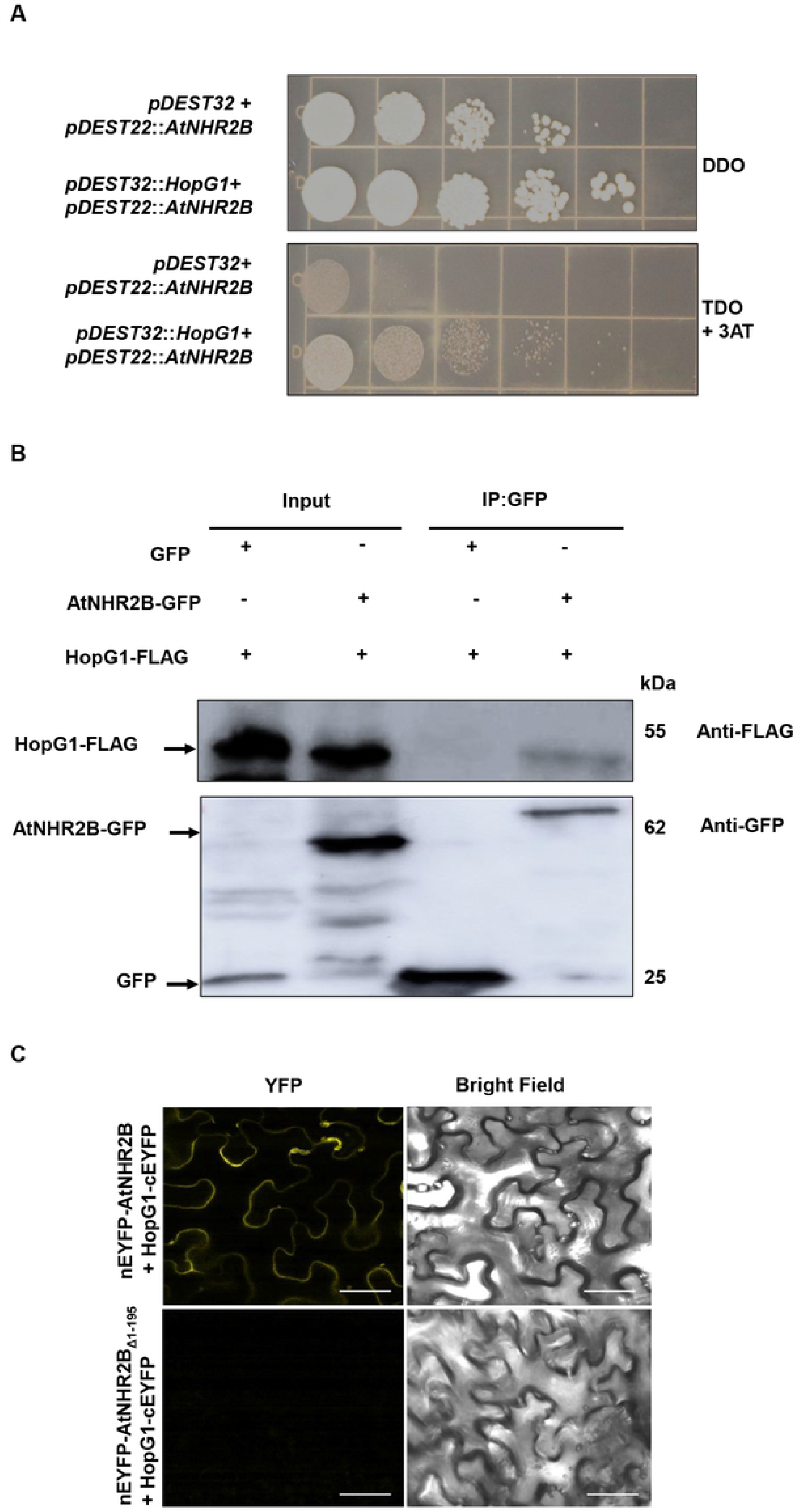
HopG1 interacts with AtNHR2B. (**A**) HopG1 interacts with AtNHR2B in yeast. Yeast strain MaV203 was co-transformed with *pDEST32::HopG1* and *pDEST22:: AtNHR2B*, or *pDEST32* and *pDEST22::AtNHR2B* and transformants were isolated in on Double Drop Out (DDO, SD/-Leu/-Trp) media and transferred to Triple Drop Out (TDO, SD/-His/-Leu/-Trp) + 15 mM 3-Amino-1,2,4-Triazole. All the experiments were repeated three times and shown similar results. (**B**) HopG1 and AtNHR2B-GFP interact in planta. *HopG1-FLAG* and *AtNHR2B-GFP*, or *HopG1-FLAG* and *35S-GFP* combinations were transiently co-expressed in five-week-old *N. benthamiana* plants. Infiltrated leaves were harvested for protein extraction followed by immunoprecipitation using GFP Nanobody/VHH coupled to agarose beads. Immunoprecipitated samples were separated by SDS-PAGE electrophoresis and transferred to a Polyvinylidene difluoride (PVDF) membrane for Western Blot analysis using anti-FLAG and anti-GFP antibodies. (**C**) Bimolecular fluorescence complementation analysis shows the interaction between HopG1 and AtNHR2B in *planta. HopG1* fused to the C-terminal half of *EYFP* (*HopG1-cEYFP*), was transiently co-expressed in *N. benthamiana* with *AtNHR2B* fused to the N-terminal half of *EYFP* (*nEYFP-AtNHR2B*), or with the truncated version AtNHR2B _Δ1-195_ fused to the N-terminal half (*nEFYP*-AtNHR2B _Δ1-195_). At 3 dpi, infiltrated leaves were imaged by laser scanning confocal microscopy using excitation and emission wavelengths of 488 nm and 500 to 535 nm, respectively. Confocal images were taken at 63X magnification. Scale bars = 50 μm.

To evaluate the HopG1/AtNHR2B interaction in the appropriate biological context, we transiently co-expressed HopG1 fused to the flag epitope (HopG1-FLAG) with AtNHR2B-GFP for co-immunoprecipitation. HopG1-FLAG and free GFP were also co-infiltrated and used as control. Immunoprecipitation of AtNHR2B-GFP using GFP Nanobody/VHH coupled to agarose beads (GFP-Trap® Agarose, Chromotek), co-immunoprecipitated HopG1-FLAG as detected by Western Blot using Anti-Flag antibodies (Figure 3B). In contrast, immunoprecipitation of free GFP did not co-immunoprecipitate HopG1-FLAG, demonstrating that the physical interaction between HopG1 and AtNHR2B also occur in planta. We further evaluated the in-situ interaction of AtNHR2B and HopG1 by co-expressing AtNHR2B fused to the N-terminal half of the enhanced yellow fluorescent protein (nEYFP) and the HopG1 fused to the C-terminal half of EYFP (cEYFP) for Bimolecular fluorescence complementation (BiFC) assays (33). The in-situ interaction between AtNHR2B and HopG1 was revealed by the restoration of the yellow fluorescence and shown to occur in the cytoplasm; importantly, amino acids 1-195 in AtNHR2B are important for the interaction as deletion of the region encoding those amino acids abolishes the interaction with HopG1 (Figure 3C).

### HopG1 expression causes virulence-related induced cell death in *N. benthamiana* that is suppressed by AtNHR2B

To further understand how the interaction between HopG1 and AtNHR2B in planta was related with the function of HopG1, we transiently co-expressed *HopG1-FLAG* with free *GFP* or in combination with *AtNHR2B-GFP* in *N. benthamiana*. At 4 days after infiltration, the section of the leaf expressing *HopG1-FLAG* alone (Supplementary Figure 1A), or co-expressed with *GFP* showed extensive tissue necrosis but that tissue necrosis was reduced in the section of the leaf co-expressing *HopG1-FLAG* and *AtNHR2B-GFP* (Figure 4A). Similar results were obtained when transiently expressing another version of HopG1 epitope-tagged with HA (Supplementary Figure 1B). To further understand how AtNHR2B reduced tissue necrosis caused by HopG1, we evaluated the HopG1 protein abundance by Western blot. Tissues co-expressing *HopG1-FLAG* with *AtNHR2B-GFP* had reduce levels of *HopG1-FLAG* in comparison with tissues co-expressing *HopG1-FLAG* with free *GFP* (Figure 4B). Similar results were obtained when transiently expressing another version of HopG1 epitope-tagged with HA (Supplementary Figure 1C). Co-expression of *AtNHR2B-GFP* with *HopG1-FLAG* do not alter the AtNHR2B-GFP protein abundance (Supplementary Figure 2). Collectively, these results indicate that HopG1 transiently expressed in *N. benthamiana* induces cell death that is attenuated by AtNHR2B degrading HopG1.

**Figure 4.**
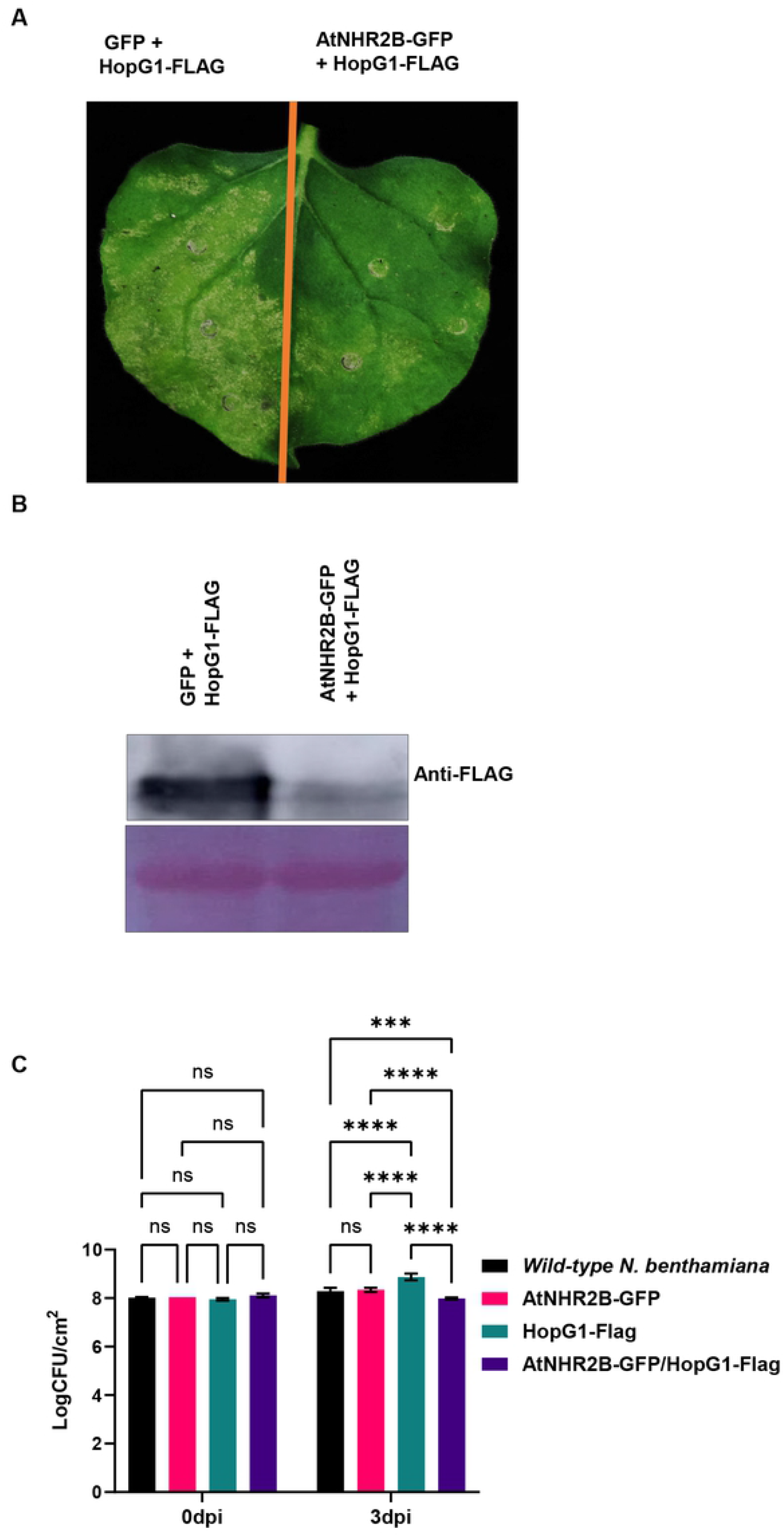
Transient expression of HopG1 in *N. benthamiana* triggers cell death that is attenuated by AtNHR2B. (**A**) Phenotypic evaluation of cell death in *N. benthamiana. HopG1-FLAG* was transiently co-expressed in *N. benthamiana* with *AtNHR2B-GFP* or with *GFP* alone. At 24 hpi, 20μM DEX was sprayed to induce the expression of *HopG1-FLAG*. Cell death was evaluated at 4 dpi. (**B**) AtNHR2B alters the abundance of HopG1 protein. *HopG1-FLAG* was transiently co-expressed in *N. benthamiana* with *AtNHR2B-GFP* or with *GFP* alone. At 24 hpi, 20μM DEX was sprayed to induce the expression of *HopG1-FLAG*. Inoculated leaves were collected 72 hpi for protein extraction and Western Blot analysis using anti-FLAG antibodies. (**C**) Transient expression of *HopG1-FLAG* enhances *Pst* DC3000 growth in *N. benthamiana*. Wild-type *N. benthamiana* and *N. benthamiana* plants transiently expressing *AtNHR2B-GFP, HopG1-FLAG* and *AtNHR2B-GFP/HopG1-FLAG*, were syringe-inoculated with the non-adapted bacterial pathogen *Pst* DC3000 at a concentration of 1 × 10^5^CFU/ml. *HopG1-FLAG* and *AtNHR2B-GFP/HopG1-FLAG* expressing leaves were sprayed with 20μM DEX 24h prior *Pst* DC3000 inoculation. Inoculated leaves were collected at 0 and 3 dpi to enumerate bacterial populations. Bars represent means and standard deviation for bacterial growth in wild-type *N. benthamiana* and *N. benthamina* plants transiently expressing *AtNHR2B-GFP, HopG1-FLAG* and *AtNHR2B-GFP/HopG1-FLAG*. Asterisks above the bars represent statistically significant differences in the growth of wild-type *Pst DC3000* in wild-type *N. benthamiana* and *N. benthamina* plants transiently expressing *AtNHR2B-GFP, HopG1-FLAG* and *AtNHR2B-GFP/HopG1-FLAG* using a two-way ANOVA and Sidak’s multiple comparison test with P-value ≤0.05. Number of asterisks reflects levels of significance, with ns equivalent to P≥0.05, * to P≤0.05, ** to P≤0.01, *** to P≤0.001 and **** to P≤0.0001. The experiments were repeated three times with consistent results.

Since the cell death phenotype can be HR or necrosis associated with virulence, we resolved between those two alternatives by inoculating *N. benthamiana* plants transiently expressing *HopG1-FLAG* with the non-adapted pathogen of *N. benthamiana PstDC3000* that does not grow in wild-type *N. benthamiana*. The results showed that *Pst* DC3000 did not grow in wild-type *N. benthamiana* nor in plants expressing *AtNHR2B-GFP*, as the bacterial populations at 3 dpi were similar to the initial populations at 0 dpi (Figure 4C). Interestingly, the populations of *Pst* DC300 in at 3 dpi in *N. benthamiana* plants expressing *HopG1-FLAG* had slightly higher populations that were statistically significant in comparison with the populations in wild-type *N. benthamiana* plants or in *N. benthamiana* plants expressing *AtNHR2B-GFP*. Furthermore, similar to the reduction in cell death phenotype and HopG1-FLAG accumulation in *N. benthamiana* plants co-expressing *AtNHR2B-GFP* with *HopG1-FLAG*, inoculation of *Pst* DC3000 in *N. benthamiana* plants co-expressing *AtNHR2B-GFP* and *HopG1-FLAG* also contributed to reduced populations of *Pst* DC3000 at 3dpi (Figure 4C). Taken together, these results show that HopG1 triggers plant cell death that is not an HR, but either contributes to plant susceptibility by enabling the growth of a non-adapted pathogen, or contributes directly to enhance virulence of *Pst* DC3000. Moreover, the virulence activities of HopG1-inducing cell death to promote pathogen multiplication, are actively attenuated by AtNHR2B.

### Arabidopsis transgenic plants expressing HopG1-FLAG induce mitochondrial ROS-related cell death that is attenuated by AtNHR2B

To further validate the results obtained in *N. benthamiana*, we obtained transgenic *Arabidopsis* lines expressing *HopG1-FLAG* under dexamethasone inducible promoter, and further crossed them with transgenic lines expressing *AtNHR2B-GFP*. Wild-type Col-0 plants and transgenic plants expressing *HopG1-FLAG, AtNHR2B-GFP* and *HopG1-FLAG/AtNHR2B-GFP* were inoculated with the non-adapted pathogen *Pstab* or mock-treated with water to evaluate cell death after staining with Trypan Blue (34). Inoculation with *Pstab* triggered cell death in transgenic plants expressing *HopG1-FLAG* alone or in combination with *AtNHR2B-GFP*, whereas, no cell death was observed in wild-type Col-0 plants nor in plants expressing *AtNHR2B-GFP* alone. Importantly, less cell death was observed in transgenic plants co-expressing *HopG1-FLAG* with *AtNHR2B-GFP*, suggesting that AtNHR2B-GFP attenuates the HopG1-triggered cell death (Figure 5A). We confirmed that to be the case by using the same plants’ genotypes and pathogen inoculation conditions to quantify cell death by electrolyte leakage at 5-, 10-, 20- and 24-hours post-inoculation (hpi). The results showed that at 20 hpi, plants expressing *HopG1-FLAG* have the highest levels of electrolyte leakage that were significantly different from the electrolyte leakage levels in the other plant genotypes, while plants expressing *AtNHR2B-GFP* had the lowest levels, and plants co-expressing *HopG1-FLAG* and *AtNHR2B-GFP* had intermediate levels. Interestingly, at 24 hpi, the levels of electrolyte leakage between plants expressing *HopG1-FLAG* alone and plants expressing *HopG1-FLAG* together with AtNHR2B-GFP are not significantly different supporting the idea that AtNHR2B is attenuating the HopG1-triggered cell death (Figure 5B).

**Figure 5.**
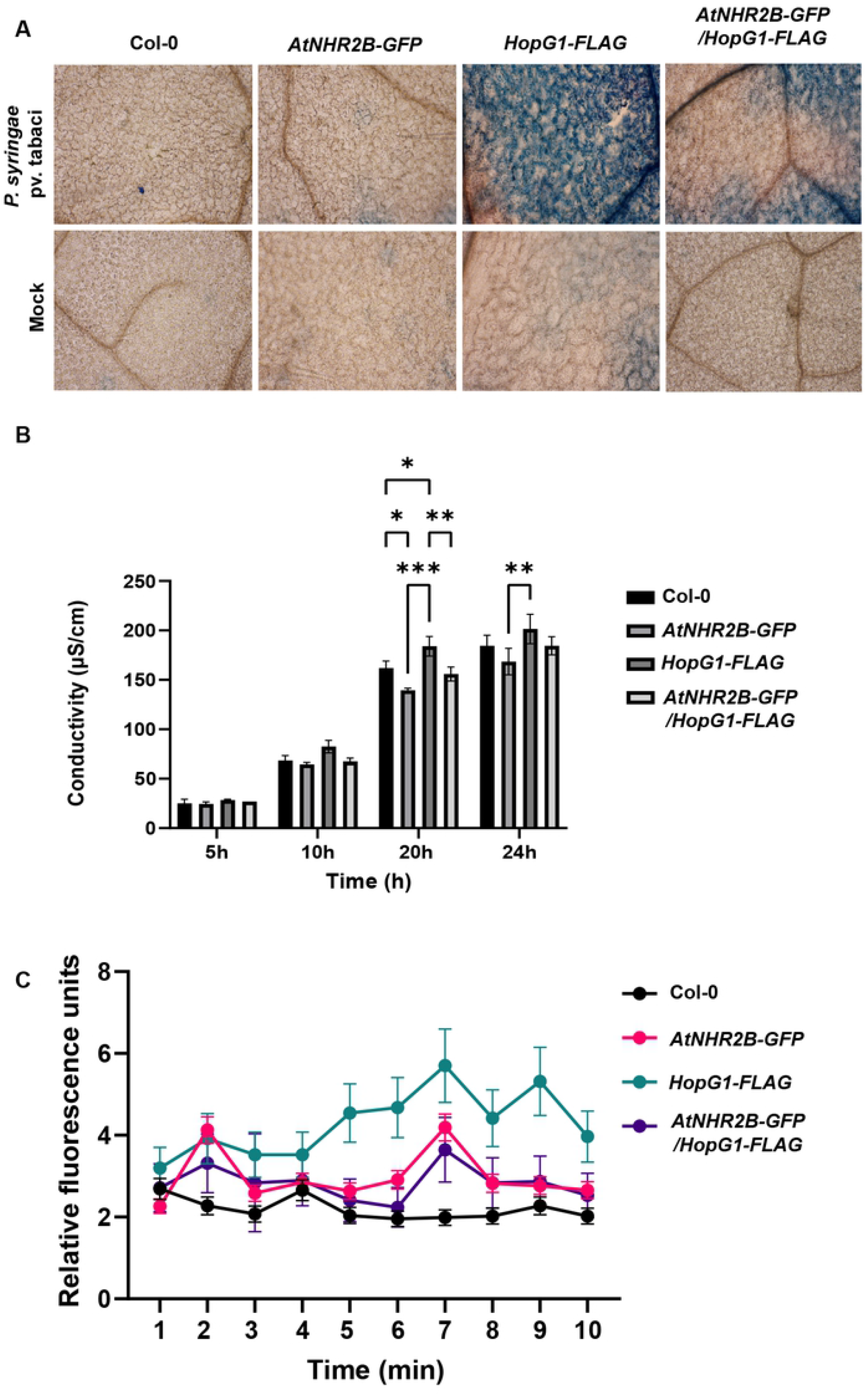
Transgenic expression of HopG1 in Arabidopsis causes cell death that is attenuated by AtNHR2B. (**A**) Cell death phenotype in *HopG1* expressing plants is reduced by expression of *AtNHR2B*. Five-week-old wild-type Col-0, *AtNHR2B-GFP, HopG1-FLAG* and *AtNHR2B-GFP/HopG1-FLAG* plants were syringe-inoculated with *Pstab* at a concentration of 1 X 10^6^ CFU/mL or infiltrated with water (mock). *HopG1-FLAG* and *AtNHR2B-GFP/HopG1-FLAG* plants were sprayed with 20μM DEX 24h prior bacterial inoculation. At 24 hpi, treated leaves were detached and stained with 0.05% trypan blue. Images were taken on a light microscope using bright field. (**B**) Ion leakage is enhanced in plants expressing *HopG1*. Five-week-old wild-type Col-0 and transgenic plants expressing *AtNHR2B-GFP, HopG1-FLAG*, and *AtNHR2B-GFP/HopG1-FLAG* were syringe-inoculated with *Pstab* at a concentration of 1 × 10^6^CFU/ml to evaluate electrolyte leakage at 5, 10, 20 and 24 hpi. *HopG1-FLAG* and *AtNHR2B-GFP/HopG1-FLAG* were sprayed with 20μM DEX 24h prior bacterial inoculation. Bars represent means of conductivity (μS/cm) and asterisks above the bars represent statistically significant difference with P-value ≤ 0.05 using a two-way ANOVA and Tukey’s multiple comparison test. Number of asterisks reflects levels of significance, with ns equivalent to P≥0.05, * to P≤0.05, ** to P≤0.01, *** to P≤0.001 and **** to P≤0.0001. (**C**) *Arabidopsis* plants expressing HopG1-FLAG generate higher levels of ROS from mitochondrial origin. Wild-type Col-0 and transgenic plants expressing *HopG1-FLAG, AtNHR2B-GFP* and *AtNHR2B-GFP/HopG1-FLAG* were flood-inoculated with the non-adapted bacterial pathogen *Pstab* at a concentration of 1x 10^7^CFU/ml, or infiltrated with water (mock). *HopG1-FLAG* and *AtNHR2B-GFP/HopG1-FLAG* were sprayed with 20μM DEX 24h prior bacterial inoculation. Leaf disks were incubated with MitoTracker Red CM-H_2_XRos and mitochondrial ROS fluorescence was measured using excitation/emission wavelengths of 570 nm and 535 nm, respectively.

Because cell death phenotypes are regulated by reactive oxygen species (ROS), and mitochondria are the source of ROS, it was necessary to evaluate how HopG1 contributes to the production of ROS in the mitochondria. For that purpose, a mitochondria-specific ROS sensor, MitoTracker Red CM-H_2_XRos (ThermoFisher Scientific, Waltham, MA) was used to evaluate mitochondrial ROS produced after pathogen infection. Inoculation of *Pstab* triggered an accumulation of mitochondrial ROS in all the plants. However, the levels of accumulation varied between genotypes. The lowest levels of mitochondrial ROS were observed in wild-type Col-0, moderate levels were found in transgenic plants expressing *AtNHR2B-GFP* and co-expressing *HopG1-FLAG* and *AtNH2B-GFP*. The highest levels of mitochondrial ROS were observed in plants expressing *HopG1-Flag* alone (Figure 5C).

### HopG1 Interferes with AtNHR2B Function to Promote Disease

Our results demonstrating an interaction between HopG1 and AtNHR2B and a possible interplay between both proteins, prompted us to further investigate the antagonism between HopG1 and AtNHR2B. HopG1 was previously shown to interfere with callose deposition when *HopG1-HA* expressing plants were inoculated with the *Pst* DC3000 *hrcC* mutant (24). In our assays, we used *Pstab* that also triggers PTI in Arabidopsis as demonstrated by the callose deposits in the wild-type Col-0 plants and transgenic Arabidopsis plants expressing *AtNHR2B-GFP* (Figure 6A). As previously reported, transgenic plants expressing *HopG1-FLAG* were devoid of callose deposits and were comparable to mock-treated plants. Similarly, plants co-expressing *HopG1-FLAG* and *AtNHR2B-GFP* were also devoid of callose deposits suggesting that AtNHR2B is unable to interfere with the PTI-suppressing activities of HopG1 (Figure 6A).

**Figure 6.**
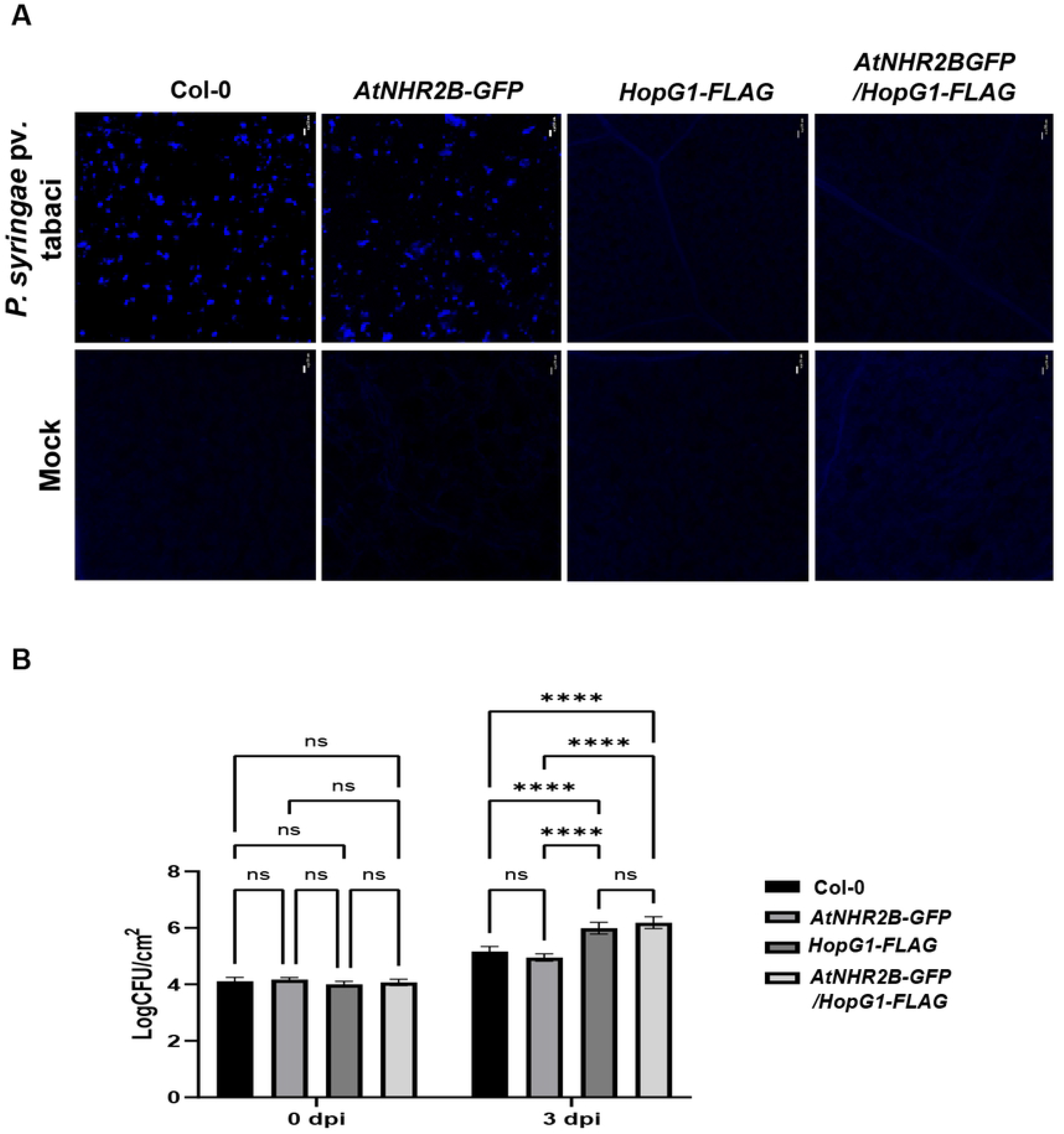
Transgenic expression of HopG1 interferes with AtNHR2B-dependent plant defense responses. (**A**) Plants expressing *HopG1* are deficient in callose deposition. Five-week-old wild-type Col-0, *AtNHR2B-GFP, HopG1-FLAG*, and *AtNHR2B-GFP/HopG1-FLAG* were syringe-inoculated with the non-adapted bacterial pathogen *Pstab* at a concentration of 1 × 10^7^CFU/ml. *HopG1-FLAG* and *AtNHR2B-GFP*/*HopG1-FLAG* were sprayed with 20μM DEX 24h prior bacterial inoculation. Inoculated leaves were detached at 24 hpi and stained with 5% aniline blue staining to evaluate callose deposition. Images were taken using a confocal microscope under DAPI filter. Scale bar = 20 μm. (**B**) Transgenic expression of HopG1-contributes to bacterial growth. Wild-type Col-0, *AtNHR2B-GFP, HopG1-FLAG*, and *AtNHR2B-GFP/HopG1-FLAG* were syringe inoculated with *Pstab* at a concentration of 1x 10^6^CFU/ml. *HopG1-FLAG* and *AtNHR2B-GFP*/*HopG1-FLAG* were sprayed with 20μM DEX 24h prior bacterial inoculation. Leaf samples were collected at 0 and 3 dpi. Bars represent means of CFU/cm^2^. Asterisks above the bars represent statistically significant difference with P-value ≤ 0.05 using a two-way ANOVA and Sidak’s multiple comparison test. Number of asterisks reflects levels of significance, with ns equivalent to P≥0.05, * to P≤0.05, ** to P≤0.01, *** to P≤0.001 and **** to P≤0.0001. All above experiments were repeated three times with similar results.

Consistent with the activities of HopG1 suppressing PTI and the inability of AtNHR2B to interfere with HopG1-mediated PTI suppression, transgenic plants expressing *HopG1-FLAG* alone or in combination with AtNHR2B-GFP supported 10-fold higher growth of *Pstab* at 3dpi populations in comparison with *Pstab* growth in Col-0 or plants expressing *AtNHR2B-GFP* (Figure 6B). Taken together, these results demonstrate that HopG1 interferes with plant defense responses even when AtNHR2B-GFP is overexpressed. Moreover, the increased bacterial growth and lack of callose deposition, suggest that similar to the results obtained in *N. benthamiana*, the HopG1-mediated cell death are not related to the HR but to a virulence mechanism related to HopG1 function.

## Discussion

*Pst* DC3000 deploys a plethora of effectors into the plant cell to interfere with plant defense responses, alter cellular processes and promote bacterial parasitism (6, 7). The *Pst* DC3000 T3E HopG1 appears to be of paramount importance in the pathogenicity of *Pst* DC3000 by being one of five effectors constituting the minimal repertoire that makes *Pst DC3000*Δ*hopQ1-1* pathogenic in *N. benthamiana* (4, 23). Previous studies also demonstrated that HopG1 suppresses the HR in *N. benthamiana*, as only the *Pst* DC3000 *hopG1* mutant, but not with the (26). Those results suggest that the absenceSimilarly, a *Pst* DC3000 strain deleted of all effectors and only harboring *HopG1* failed to elicit the HR in *N. benthamiana* (35). Furthermore, transient expression of the cell death inducer *BAX1* (36) in *N. benthamiana* triggered cell death as expected, but this cell death was not observed when *BAX1* was transiently co-expressed with *HopG1* (26).

In addition to suppressing the HR, transgenic expression of *HopG1* fused to the haemagglutinin (HA) epitope in Arabidopsis also led to suppression of PTI, observed as a reduction in callose deposition after infiltration with the Flg 21 peptide, or after inoculation with the non-pathogen *Pseudomonas fluorescens* (24). We confirmed these results by showing that Arabidopsis plants expressing *HopG1-FLAG* and inoculated with the non-adapted pathogen *Pstab* are also devoid of callose deposits.

Other previous observations on the virulence function of HopG1, revealed its role in the remodeling of the cytoskeleton demonstrated to occur through the interaction of HopG1 with the mitochondrial-localized kinesin motor protein (27). The mitochondrial localization of HopG1 previously led to the hypothesis that HopG1 alters mitochondrial function, and in support of that hypothesis, Arabidopsis plants expressing *HopG1-HA* showed reduced oxygen consumption and enhanced ROS levels (24). However, that study used the ROS sensitive probe H2DCFDA (2’-7’-dichlorodihydrofluorescein), that does not discriminate among the multiple sources of ROS. In this study, we use a mitochondria-specific fluorogenic ROS sensor and defined more precisely that the higher levels of ROS in *HopG1*-FLAG transgenic lines are from mitochondrial origin. Thus, our results provide stronger evidence that *HopG1-FLAG* expression and localization to mitochondria actually induces the production of mitochondrial ROS that likely activate a cell death program.

In this study, we uncovered that indeed HopG1 induces necrotic cell death by transient expression in *N. benthamiana* and by stable expression in Arabidopsis. Previously, *Agrobacterium-mediated* transient expression of *Pst* DC3000 HopG1 in 59 plant accessions that included *N. benthamiana*, revealed that this effector caused cell death in a limited number of plants (ca 5) (37). Such cell death phenotype was explained as a recognition by the plant (HR) given that the effector was from a non-adapted pathogen (37). Our data demonstrate that actually, the cell-death phenotype observed is the result of the virulence function of HopG1.

The cell death phenotypes that effectors trigger can be difficult to assess as they might correspond to completely opposite outcomes; the HR, a plant defense response that restricts pathogen proliferation, or a pathogen-induced necrosis that sustains pathogen proliferation. Thus, to resolve if the HopG1-dependent cell death was HR or virulence-related necrosis, we inoculated the *HopG1-FLAG*-expressing *N. benthamiana* plants *with* the non-adapted pathogen *Pst* DC3000. The higher levels of *Pst* DC3000 populations in *HopG1*-expressing plants in comparison with plants not expressing *HopG1* indicated that the observed cell death was a necrotic-type of cell death that enables *Pst* DC3000 to proliferate to higher numbers in a nonhost plant. Similarly, transgenic Arabidopsis plants expressing *HopG1-FLAG* also had enhanced cell death that enabled the growth of the non-adapted pathogen of Arabidopsis, *Pstab*. This necrotic cell death agrees with previous results showing that HopG1 is required for the development of chlorotic symptoms (27).

We also uncovered a functional relationship between HopG1 and AtNHR2B based on the following lines of evidence: 1) *Atnhr2b* mutation restores the growth deficiency of the *Pst* DC3000 Δ*hopG1*; 2) HopG1 directly interacts with AtNHR2B in yeast and in planta; 3) AtNHR2B suppresses the necrotic cell death induced by HopG1. The results suggest that HopG1 and AtNHR2B counteract each other with HopG1 inducing necrotic cell death and AtNHR2B attenuating HopG1-induced cell death. Moreover, in *N. benthamiana*, expression of *HopG1-FLAG* enables the growth of the non-adapted pathogen *Pst* DC3000, whereas co-expression of *HopG1-FLAG* and *AtNHR2B-GFP* in *N. benthamiana* effectively reduces *Pst* DC3000 populations. Interestingly, HopG1-induced callose deposition in Arabidopsis is not attenuated by AtNHR2B and *Pstab* bacterial populations are the same in plants expressing *HopG1-FLAG* alone or co-expressing *HopG1-FLAG* and *AtNHR2B-GFP*. Altogether, these results would indicate a specific function of AtNHR2B attenuating necrotic cell death only through targeted degradation of HopG1.

The enhanced growth of *Pstab* in Arabidopsis plants expressing *HopG1-FLAG* alone and also in plants co-expressing *HopG1-FLAG* and *AtNHR2B-GFP* can also indicate that transgenic expression of *HopG1-FLAG* enhances the virulence of *Pstab* as *Pstab* do not encode *HopG1* (38). Thus, given the importance of HopG1 in the pathogenesis of *Pst* DC3000, it is also possible that HopG1 augments the virulence of *Pstab* hampering with AtNHR2B ability to overcome it.

Collectively, our results and the results of others highlight that HopG1 is a multi-faceted protein that can suppress the HR, interfere with plant defense responses but also cause necrotic cell death associated with chlorotic symptoms. All these phenotypes associated with HopG1 highlight is function early in infection, suppressing immune responses, and later in infection triggering necrotic cell death. The functions of HopG1 are consistent with the hemi-biotrophic lifestyle of *Pst* DC3000 that combines an early biotrophic phase suppressing cell death responses with a late necrotrophic phase inducing cell death.

## Materials and Methods

### Bacterial Strains

Wild-type *P. syringae* pv tabaci (*Pstab*), *Pst* DC3000 and *Pst* DC3000Δ*hopG1* mutant were grown on King’s B (KB) medium supplemented with rifampicin (25 μg/mL). *Agrobacterium tumefaciens* strains were grown in Luria-Bertani (LB) medium supplemented with rifampicin (25 μg/mL) and kanamycin (50 μg/ml). All the strains were grown at 28°C. Bacterial strains used in this study are listed in Table S1.

### Plant Materials and Growth Conditions

*Arabidopsis thaliana* seeds were planted in soil for two weeks, transplanted to individual pots and grown for four more weeks. Plants were grown in growth chambers at 21°C with an 8/16 h light/dark cycles. *Nicotiana benthamiana* seeds were sown in soil, transplanted after 2 weeks and grown for 4 weeks under growth chamber conditions at 25°C with a 10/14 h light/dark cycle.

Transgenic lines expressing *HopG1-FLAG* under the expression of the glucocorticoid promoter were obtained from Dr. Jim Alfano (University of Nebraska, Lincoln). *HopG1-FLAG* plants were crossed with lines expressing *AtNHR2B-GFP* to generate *AtNHR2B-GFP/HopG1-FLAG* transgenic lines.

### Plasmid constructs

pENTR/SD:HopG1 (35) was transferred into the yeast bait vector *pDEST32* through an LR reaction of Gateway Cloning (Thermo Fisher Scientific, Waltham, MA) to generate a transcriptional fusion to the GAL4-DNA binding domain. *AtNHR2B* in the entry vector *pDONR201* (19) was cloned into the yeast prey vector *pDEST22* to generate a fusion to the GAL4 activation domain.

### Transient expression in *N. benthamiana*

*Agrobacterium tumefaciens* GV2260 harboring constructs of interest were induced and infiltrated into fully expanded leaves of 3-week old *N. benthamiana* plants using a needle-less syringe as previously described (31). Infiltrated leaves were used to evaluate protein localization or in situ protein-protein interaction by laser scanning fluorescence microscopy, or to evaluate protein expression or evaluate protein-protein interaction by co-immunoprecipitation followed by Western blot.

### Western blot

*N. benthamiana* transiently expressing proteins of interest were collected in liquid nitrogen. Tissue was ground in liquid nitrogen and homogenized with protein extraction buffer (50 mM Hepes-pH 7.5, 250 mM sucrose, 10 mM EDTA, 5% glycerol, 50 mM sodium pyrophosphate, 25 mM sodium fluoride, 1 mM sodium molybdate, 3 Mm DTT, 1 mM PMSF and plant protease inhibitor cocktail) (Thermo Scientific) (Heese et al., 2007). *Arabidopsis* plants expressing *AtNHR2B-GFP* were harvested in liquid nitrogen and homogenized with protein extraction buffer (100 mM Tris-HCl, pH 7.5, 150 mM NaCl, 1 mM EDTA, 10 mM MgCl_2_ 0.20% NP40, 0.1% SDS, 5 mM DTT, 10% glycerol, 1 mM phenylmethylsulfonyl fluoride (PMSF) and plant protease inhibitor cocktail) (Thermo Scientific).

The concentrations of protein in the supernatant were determined by using the Bio-Rad protein assay reagent (Bio-Rad). Samples, typically 30-50 mg, were separated on 10 % acrylamide containing SDS-PAGE gels (mini protean; Bio-Rad) and transferred to polyvinylidene fluoride membrane (GE Healthcare). Membranes were incubated with appropriate antibodies: anti-GFP (1:2,000 dilution; Invitrogen), Anti-FLAG (A5892, Sigma) and anti-HA antibody (H6908, Sigma), were used at dilution of 1:1,000. Rabbit antisera (GE Healthcare) was used as secondary antibody with 1:5,000 dilution. Chemiluminescent detection was done by using the ImageQuant™ LS-500 image system (GE Healthcare).

### Live-cell imaging

*N. benthamiana* plants transiently expressing *AtNHR2B-GFP*, co-expressing *AtNHR2B-nEYFP* with *HopG1-cEYFP*, or *AtNHR2BΔ_1-195n_EYP with HopG1-cEYFP* were used to evaluate *AtNHR2B-GFP* localization, or *AtNHR2B/HopG1* interaction by bimolecular fluorescence complementation (BiFC), respectively. GFP and YFP signals were imaged on a Leica SP2 or Leica Stellaris 8 Laser Scanning Confocal Microscope (Leica Microsystems, Buffalo Grove, IL) at 48hpi. Arabidopsis plants expressing *AtNHR2B-GFP* were also evaluated by laser scanning fluorescence microscopy. GFP and YFP fluorescence was imaged using excitation wavelength of 488 nm and an emission wavelength of 500 to 535 nm.

### Bacterial Inoculation into *Arabidopsis thaliana* plants

Five-week-old *Arabidopsis thaliana* plants, genotypes wild-type Col-0, *Atnhr2b* mutant, or expressing *AtNHR2B-GFP, HopG1-FLAG* and *AtNHR2B-GFP/HopG1-FLAG* were syringe-inoculated with *PstDC3000, PstDC3000 (ΔhopG1*) or *Pstab* at different concentrations depending on the experiment.

### Bacterial multiplication assays

Leaf disks (0.5 cm^2^) from inoculated plants were collected, serially-diluted and plated as previously described (31). Each experiment was repeated three times.

### Yeast two-hybrid assay

The yeast strain Mav203 was grown in YPDA at 30°C overnight with constant shaking. The OD_600_ of overnight grown cultures was measured, diluted to an OD_600_ of 0.4 in a final volume of 50 ml of YPAD and grown for additional 3h. Yeast cells in mid-log phase were co-transformed with pDEST32/pDEST22::*AtNHR2B* or pDEST32::*HopG1/* pDEST22::*AtNHR2B* using the Frozen-EZ Yeast Transformation II Kit (Zymo Research, Irvine, CA). Transformed yeast cells were plated on Double Drop Out (DDO, SD/-Leu/-Trp) selection plates and grown at 30°C for 4 days. Single colonies were picked from the plates and cultured in 15 mL DDO broth at 30°C overnight. The overnight culture was diluted to an OD_600_ of 0.2 and plated on triple dropout medium (TDO, SD/-His/-Leu/-Trp) containing 15 mM 3-Amino-1,2,4-Triazole (3-AT) and grown at 30°C for 4 days.

### Co-immunoprecipitation

Four-week-old *N. benthamiana* plants were co-infiltrated with two strains of *A. tumefaciens* strains harboring *AtNHR2B-GFP* and *HopG1-Flag*, or *A. tumefaciens* strains harboring *AtNHR2B-GFP* and *35S-GFP*. Infiltrated leaves were collected at 2 days post-inoculation and tissue, ground in liquid nitrogen and homogenized in 1 mL of co-immunoprecipitation extraction buffer (100 mM Tris-HCl, pH 7.5, 150 mM NaCl, 1 mM EDTA, 10 mM MgCl2, 10% Glycerol, 0.2% Nonidet P-40, 1 mM PMSF, 5 mM DTT, 1X Proteinase inhibitor cocktail (Sigma Aldrich, St. Louis, MO). Protein extracts were incubated for 30 min on ice and centrifuged at 4°C for 30 min at 13,000 rpm. Supernatants containing extracted proteins were collected in a pre-chilled 1mL Eppendorf tube. Protein concentration was measured by Bradford Assay (BioRad, Hercules, CA) and protein expression in input samples was confirmed by Western blot as described above.

One milligram of total protein extract was mixed with 25 μl of GFP Nanobody/VHH coupled to agarose beads (GFP-Trap® Agarose, Chromotek) and incubated for 3 hours and 30 minutes at 4°C with end to end rocking. After incubation, protein complexes bound to beads were washed once with 1x TBS buffer (50 mM Tris-HCl, 150 mM NaCl, pH 7.5) and twice with 1x TBS buffer (50 mM Tris-HCl, 500 mM NaCl, pH 7.5). Protein complexes bound to the beads were eluted in 2x SDS protein loading buffer, loaded and ran into an SDS-PAGE gel and transferred to PVDF membranes. Proteins were detected by Western Blot as described above.

### Cell death assay

Five-week old wild-type Col-0, *AtNHR2B-GFP, HopG1-FLAG* and *AtNHR2B-GFP/HopG1-FLAG* plants were syringe-inoculated with *Pstab* at OD_600_=0.02 (1 X 10^6^ CFU/mL). Control plants were inoculated with water only. At 24 hpi, six to nine inoculated leaves were detached from six independent plants for each genotype. Collected leaves were stained with 0.05% trypan blue for 45 minutes at room temperature and washed twice with PBS (34). Images were taken on a light microscope using bright field.

### Callose Deposition

Five-week old wild-type Col-0, *AtNHR2B-GFP, HopG1-FLAG* and *HopG1-FLAG/AtNHR2B-GFP* were sprayed with 20 μM dexamethasone to induce expression of *HopG1-FLAG*. At 24 h after dexamethasone treatment, plants were syringe-inoculated with *Pstab* at OD_600_=0.02 (1 X 10^6^ CFU/ml). Ten leaves from six independent plants for each genotype and inoculated with *Pstab* or infiltrated with water were detached after 24 hpi and stained with 5% aniline blue to visualize callose deposits (39). Images were taken by Nikon 90i upright scanning laser confocal microscope (Nikon) using a DAPI (4’,6-diamidino-2-phenylindole) filter with excitation wavelength of 405 nm and an emission wavelength of 450-510 nm.

### Electrolyte leakage assay

Five-week old wild-type Col-0, *AtNHR2B-GFP, HopG1-FLAG* and *AtNHR2B-GFP/HopG1-FLAG* plants were induced by dexamethasone prior to inoculation with *Pstab* at OD_600_=0.02 (1 X 10^6^ CFU/ml). Twelve leaves were collected immediately after infiltration and 0.5 cm^2^ leaf disks were excise and rinsed with 50 ml for 30 min, to wash ions released during disc excision. After 30 minutes, water was replaced with 12 ml of fresh deionized water and conductivity was measured at 5, 10, 10 and 24 hpi using the Orion Star A215 conductivity cell (013005MD) (Thermo Fisher Scientific).

### Mitochondrial ROS Production

Five-week old wild-type Col-0, *AtNHR2B-GFP, HopG1-FLAG* and *AtNHR2B-GFP*/*HopG1-FLAG* plants grown on soil were sprayed with a 20 μM dexamethasone solution supplemented with 0.01% silwet, 12 hours before inoculation. Leaf disks cut out using a 1.2 cm^2^ core-borer and harvested tissue was transferred to clear bottom plates for fluorometric analysis and submerged in *Pstab* inoculum at a final concentration of 1 X 10^7^ CFU/ml. For mock-treatment, plants were submerged in water. Two hours after inoculation with either *Pstab* mock-treatment, MitoTracker Red CM-H_2_XRos (ThermoFisher Scientific, Waltham, MA) was added at a final concentration of 0.005 mM and incubated for ten minutes before taking the first reading. Fluorescence was measured with an excitation wavelength of 570 nm and an emission wavelength of 535 nm on BioTek luminescence microplate reader.

## Acknowledgements

We thank Gitta Coaker and James M. Elmore for providing the HopG1-FLAG and HopG1-HA constructs, Jim Alfano for providing the HopG1-FLAG seeds provided and Brian Kvitko for providing the *Pst* DC3000 Δ*hopG1* mutant.

## Author Contributions

Conceived and designed the experiments: CMR, RS, CRP. Performed the experiments: CRP, RS, RC, PRL. Analyzed the data: CMR, CRP, RS, RC. Wrote the paper: CMR, CRP.

## Notes

Funding: This work was supported by the National Science Foundation CAREER award grant (IOS # 1842970) and by Equipment Grants Program, award #2021-05017, from the U. S. Department of Agriculture, National Institute of Food and Agriculture.

### Competing Interest Statement

The authors have declared no competing interest.

## References

1. Xin XF, Kvitko B, He SY. Pseudomonas syringae: what it takes to be a pathogen. Nat Rev Microbiol. 2018;16(5):316–28.

2. Xin XF, He SY. Pseudomonas syringae pv. tomato DC3000: a model pathogen for probing disease susceptibility and hormone signaling in plants. Annu Rev Phytopathol. 2013;51:473–98.

3. Wei HL, Collmer A. Defining essential processes in plant pathogenesis with Pseudomonas syringae pv. tomato DC3000 disarmed polymutants and a subset of key type III effectors. Mol Plant Pathol. 2018;19(7):1779–94.

4. Wei CF, Kvitko BH, Shimizu R, Crabill E, Alfano JR, Lin NC, et al. A Pseudomonas syringae pv. tomato DC3000 mutant lacking the type III effector HopQ1-1 is able to cause disease in the model plant Nicotiana benthamiana. Plant J. 2007;51(1):32–46.

5. Collmer A, Badel JL, Charkowski AO, Deng WL, Fouts DE, Ramos AR, et al. Pseudomonas syringae Hrp type III secretion system and effector proteins. Proc Natl Acad Sci U S A. 2000;97(16):8770–7.

6. Lindeberg M, Cunnac S, Collmer A. Pseudomonas syringae type III effector repertoires: last words in endless arguments. Trends Microbiol. 2012;20(4):199–208.

7. Cunnac S, Lindeberg M, Collmer A. Pseudomonas syringae type III secretion system effectors: repertoires in search of functions. Curr Opin Microbiol. 2009;12(1):53–60.

8. Buttner D, He SY. Type III protein secretion in plant pathogenic bacteria. Plant physiology. 2009;150(4):1656–64.

9. Schechter LM, Vencato M, Jordan KL, Schneider SE, Schneider DJ, Collmer A. Multiple approaches to a complete inventory of Pseudomonas syringae pv. tomato DC3000 type III secretion system effector proteins. Mol Plant Microbe Interact. 2006;19(11):1180–92.

10. Petnicki-Ocwieja T, Schneider DJ, Tam VC, Chancey ST, Shan L, Jamir Y, et al. Genomewide identification of proteins secreted by the Hrp type III protein secretion system of Pseudomonas syringae pv. tomato DC3000. Proc Natl Acad Sci U S A. 2002;99(11):7652–7.

11. Lindeberg M, Cartinhour S, Myers CR, Schechter LM, Schneider DJ, Collmer A. Closing the circle on the discovery of genes encoding Hrp regulon members and type III secretion system effectors in the genomes of three model Pseudomonas syringae strains. Molecular plant-microbe interactions : MPMI. 2006;19(11):1151–8.

12. Toruno TY, Stergiopoulos I, Coaker G. Plant-Pathogen Effectors: Cellular Probes Interfering with Plant Defenses in Spatial and Temporal Manners. Annu Rev Phytopathol. 2016;54:419–41.

13. Macho AP. Subversion of plant cellular functions by bacterial type-III effectors: beyond suppression of immunity. New Phytol. 2016;210(1):51–7.

14. Wilton M, Desveaux D. Lessons learned from type III effector transgenic plants. Plant Signal Behav. 2010;5(6):746–8.

15. Jones JD, Dangl JL. The plant immune system. Nature. 2006;444(7117):323–9.

16. Chisholm ST, Coaker G, Day B, Staskawicz BJ. Host-microbe interactions: shaping the evolution of the plant immune response. Cell. 2006;124(4):803–14.

17. Zipfel C, Robatzek S, Navarro L, Oakeley EJ, Jones JD, Felix G, et al. Bacterial disease resistance in Arabidopsis through flagellin perception. Nature. 2004;428(6984):764–7.

18. Cui H, Tsuda K, Parker JE. Effector-triggered immunity: from pathogen perception to robust defense. Annual review of plant biology. 2015;66:487–511.

19. Van Doorn W, Beers E, Dangl J, Franklin-Tong V, Gallois P, Hara-Nishimura I, et al. Morphological classification of plant cell deaths. Cell Death & Differentiation. 2011;18(8):1241–6.

20. Dangl JL, Jones JD. Plant pathogens and integrated defence responses to infection. Nature. 2001;411(6839):826–33.

21. Van Der Biezen EA, Jones JD. Plant disease-resistance proteins and the gene-for-gene concept. Trends in biochemical sciences. 1998;23(12):454–6.

22. Kvitko BH, Park DH, Velasquez AC, Wei CF, Russell AB, Martin GB, et al. Deletions in the repertoire of Pseudomonas syringae pv. tomato DC3000 type III secretion effector genes reveal functional overlap among effectors. PLoS Pathog. 2009;5(4):e1000388.

23. Cunnac S, Chakravarthy S, Kvitko BH, Russell AB, Martin GB, Collmer A. Genetic disassembly and combinatorial reassembly identify a minimal functional repertoire of type III effectors in Pseudomonas syringae. Proc Natl Acad Sci U S A. 2011;108(7):2975–80.

24. Block A, Guo M, Li G, Elowsky C, Clemente TE, Alfano JR. The Pseudomonas syringae type III effector HopG1 targets mitochondria, alters plant development and suppresses plant innate immunity. Cellular microbiology. 2010;12(3):318–30.

25. Guo M, Tian F, Wamboldt Y, Alfano JR. The Majority of the Type III Effector Inventory of Pseudomonas syringae pv. tomato DC3000 Can Suppress Plant Immunity. Mol Plant Microbe Interact. 2009;22(9):1069–80.

26. Jamir Y, Guo M, Oh HS, Petnicki-Ocwieja T, Chen S, Tang X, et al. Identification of Pseudomonas syringae type III effectors that can suppress programmed cell death in plants and yeast. Plant J. 2004;37(4):554–65.

27. Shimono M, Lu YJ, Porter K, Kvitko BH, Henty-Ridilla J, Creason A, et al. The Pseudomonas syringae Type III Effector HopG1 Induces Actin Remodeling to Promote Symptom Development and Susceptibility during Infection. Plant Physiol. 2016;171(3):2239–55.

28. Heath MC. Nonhost resistance and nonspecific plant defenses. Curr Opin Plant Biol. 2000;3(4):315–9.

29. Panstruga R, Moscou MJ. What is the Molecular Basis of Nonhost Resistance? Mol Plant Microbe Interact. 2020;33(11):1253–64.

30. Senthil-Kumar M, Mysore KS. Nonhost resistance against bacterial pathogens: retrospectives and prospects. Annu Rev Phytopathol. 2013;51:407–27.

31. Singh R, Lee S, Ortega L, Ramu VS, Senthil-Kumar M, Blancaflor EB, et al. Two Chloroplast-Localized Proteins: AtNHR2A and AtNHR2B, Contribute to Callose Deposition During Nonhost Disease Resistance in Arabidopsis. Mol Plant Microbe Interact. 2018;31(12):1280–90.

32. Singh R, Liyanage R, Gupta C, Lay JO, Jr., Pereira A, Rojas CM. The Arabidopsis Proteins AtNHR2A and AtNHR2B Are Multi-Functional Proteins Integrating Plant Immunity With Other Biological Processes. Front Plant Sci. 2020; 11:232.

33. Martin K, Kopperud K, Chakrabarty R, Banerjee R, Brooks R, Goodin MM. Transient expression in Nicotiana benthamiana fluorescent marker lines provides enhanced definition of protein localization, movement and interactions in planta. Plant J. 2009;59(1):150–62.

34. Kabbage M, Williams B, Dickman MB. Cell Death Control: The Interplay of Apoptosis and Autophagy in the Pathogenicity of Sclerotinia sclerotiorum. PLoS Pathog. 2013;9(4).

35. Wei HL, Zhang W, Collmer A. Modular Study of the Type III Effector Repertoire in Pseudomonas syringae pv. tomato DC3000 Reveals a Matrix of Effector Interplay in Pathogenesis. Cell Rep. 2018;23(6):1630–8.

36. Baek D, Nam J, Koo YD, Kim DH, Lee J, Jeong JC, et al. Bax-induced cell death of Arabidopsis is meditated through reactive oxygen-dependent and -independent processes. Plant Mol Biol. 2004;56(1):15–27.

37. Wroblewski T, Caldwell KS, Piskurewicz U, Cavanaugh KA, Xu H, Kozik A, et al. Comparative large-scale analysis of interactions between several crop species and the effector repertoires from multiple pathovars of Pseudomonas and Ralstonia. Plant Physiol. 2009;150(4): 1733–49.

38. Baltrus DA, Nishimura MT, Romanchuk A, Chang JH, Mukhtar MS, Cherkis K, et al. Dynamic evolution of pathogenicity revealed by sequencing and comparative genomics of 19 Pseudomonas syringae isolates. PLoS Pathog. 2011;7(7):e1002132.

39. Kvitko BH, Park DH, Velasquez AC, Wei CF, Russell AB, Martin GB, et al. Deletions in the Repertoire of Pseudomonas syringae pv. tomato DC3000 Type III Secretion Effector Genes Reveal Functional Overlap among Effectors. PLoS Pathog. 2009;5(4):16.

